# Working memory limitations and dopamine modulation in probabilistic reasoning

**DOI:** 10.64898/2026.08.12.744547

**Authors:** Cina Aghamohammadi, Jochem van Kempen, Molly Stapleton, Alwin Gieselmann, Christopher Langdon, Alexander Thiele, Tatiana A. Engel

## Abstract

Difficult decisions require gathering evidence over extended periods, placing demands on working memory. Yet, how working memory limitations affect decision-making remains largely unknown. We trained two macaque monkeys to perform a probabilistic reasoning task that involved extended sequential evidence sampling, requiring reliance on working memory. Monkeys made choices informed by a stream of briefly presented cues, each providing probabilistic evidence about which choice would be rewarded. In both animals, choices were significantly affected by working memory decay, primacy, recency, and priming. Despite individual differences in working memory limitations, both monkeys adopted sampling strategies that made their behavior nearly optimal. To test how dopamine affects working memory constraints on evidence accumulation, we systemically applied dopamine D1 receptor agonist and antagonist drugs midway during selected sessions. Activation of D1 receptors reduced priming. Blockade of D1 receptors reduced working memory decay and the subjective evidence weights assigned to individual cues. Our results reveal that complex decisions are constrained by working memory limitations and identify dopamine as a key modulator of this process, with potential implications for cognitive disorders and their treatment.

---

Complex decisions require gathering multiple consecutive pieces of information over periods that may range from seconds to days. This evidence must be integrated and transformed into a single decision. If evidence is available only temporarily, working memory plays a vital role in decision-making, allowing the brain to use information that is no longer present in the environment to guide the choice. However, working memory is not perfect, and its limitations may impact complex decision-making^1^.

At the behavioral level, working memory is characterized by limited capacity^2–4^, temporal decay^5–7^, serial-position effects, including primacy^8, 9^ and recency^10, 11^, and choice-history effects, including priming biases toward repeating choices^12, 13^. These limitations may arise from finite neural resources and biophysical constraints at the circuit level. Interference between items competing for neural resources contributes to decay, primacy and recency^14, 15^. Biophysical constraints arise from the short membrane and synaptic time constants of single neurons, which range from a few tens to hundreds of milliseconds, whereas working memory operates on much longer timescales. Recurrent connections among neurons can extend the effective timescales in the circuit to support working memory function^16^. However, even slight deviations from optimal synaptic connectivity lead to temporal decay in working memory^17^. Additionally, residual activity from previous trials can influence initial circuit conditions leading to history-dependent biases^18–20^.

The function of working memory circuits is prominently affected by dopamine^21–25^. Dopamine affects working memory through multiple cellular mechanisms, including adjusting synaptic strength^26^, synaptic plasticity^27, 28^, neuronal gain^29, 30^, and synaptic time constants^31^. These adjustments enable dopamine to mediate value-based learning and decision-making during cognitively demanding tasks^32, 33^, primarily through D1 receptors (D1Rs) in the prefrontal cortex (PFC)^34, 35^. D1R activation potentiates ‘memory fields’ of dorsolateral prefrontal cortex (DLPFC) neurons^34–36^, which are essential for maintaining information across delays. Consequently, D1R blockade in DLPFC results in impaired performance in memory guided saccade tasks in nonhuman primates^37^.

We examined how working memory limitations, and their dependence on dopamine, affect complex decision-making behavior. We trained two monkeys to perform a sequential probabilistic reasoning task^38–40^, a variant of the weather prediction task^41^, which requires subjects to make decisions by integrating probabilistic evidence from a series of cues, each varying in its reliability in predicting the rewarding choice target. Psychophysical modeling revealed that working memory limitations affected the accumulation of evidence and the judgment about when to stop sampling. Working memory decay, priming, recency, and primacy all significantly influenced performance, with decay emerging as the primary limiting factor. A neural circuit model fitted to the behavioral data suggested that differences in recurrent self-excitation may explain individual differences in memory decay and hence decision efficacy. We further investigated how dopaminergic challenges affect this complex decision process by administering D1R agonists and antagonists systemically midway through selected sessions and analyzing their effects on behavior. The D1R challenge influenced the encoding of instantaneous subjective evidence, as well as memory decay and priming. In response to these effects, animals adjusted their behavioral strategies to maintain their control condition performance levels. Our findings reveal how the constraints imposed by the working memory limitations impact complex decision-making processes during sequential probabilistic reasoning, with D1 receptors playing a major role in modulating these limitations.

## Results

### Monkeys perform sequential probabilistic reasoning

We trained two macaque monkeys to perform a sequential probabilistic reasoning task^39^. Monkeys chose one of two peripheral targets based on a sequence of observed shapes (Fig. 1a), while fixating centrally. Each shape provided probabilistic evidence indicating which target would yield a reward if chosen, with a unique weight assigned to each shape. At the beginning of each trial, one of the targets, right or left, was chosen randomly as the rewarded target by the computer, and then the shapes were drawn from the probability distribution corresponding to this target (Fig. 1b). Since the two distributions associated with the right and left targets differed, each shape provided probabilistic leverage, signaling that one target was more likely to yield a reward than the other. Five shapes favored the right target, and the other five favored the left (Fig. 1b). Thus, each shape *i* had a weight *w_i_*encoding its leverage towards the right target:

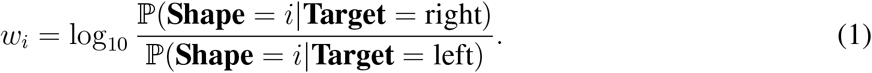

**Figure 1.**
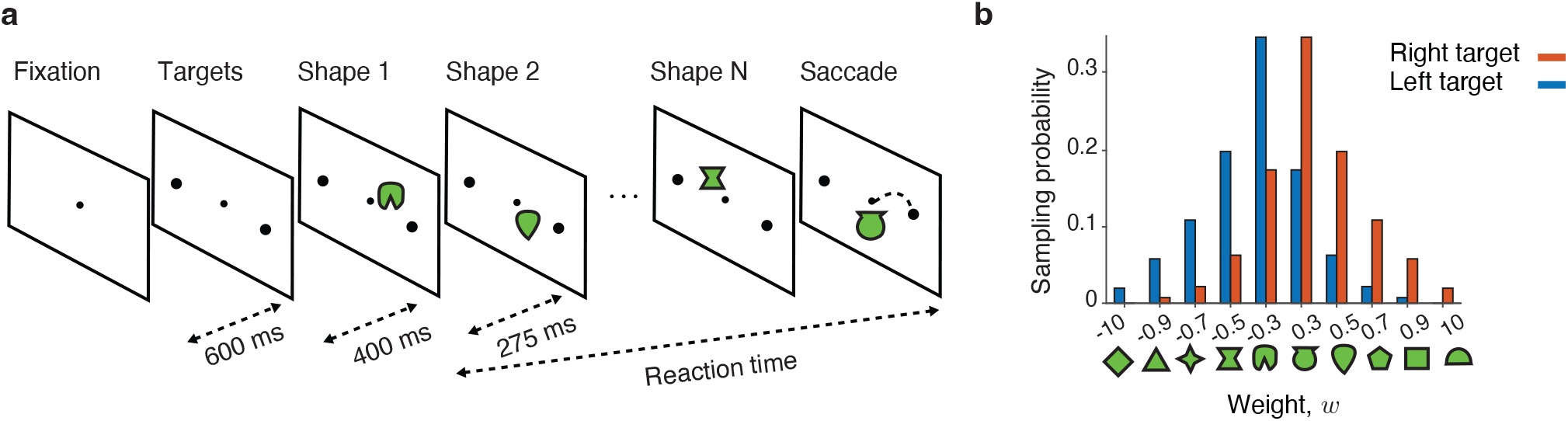
Sequential probabilistic reasoning task. **a**, A trial begins with the monkey fixating on a central point (Fixation), which must be maintained throughout the trial. After 600 ms, the right and left targets appear on the screen (Targets). The first shape appears 400 ms after target presentation, and then a new shape is presented sequentially every 275 ms. Each shape provides evidence for either the right or left choice, with varying degrees of reliability. The monkey indicates its choice with an eye movement to the selected target when ready (Saccade). The reaction time is the difference between the saccade time and the appearance of the first shape. **b**, At the start of each trial, one of the two targets, right or left, is randomly selected as rewarded target. If the right target is rewarded, the shapes are sampled from the right distribution (red); if the left target is rewarded, the shapes are sampled from the left distribution (blue).

The two shapes with the extreme weights are the trump shapes. Monkeys performed a reaction-time task, freely reporting their choice when ready. Each trial ended when the monkeys made a saccade toward the left or the right target indicating their choice, or after the presentation of 20 shapes. We collected behavioral data across multiple sessions, during which the monkeys completed a total of 81,985 trials (monkey S) and 93,065 trials (monkey T).

The monkeys gathered evidence over extended periods to guide their choices, with reaction times ranging from a few hundred milliseconds to several seconds (Fig. 2a), similar to previous studies^39^. For both monkeys, the shape displayed on the screen immediately before the saccade did not influence their choice (Supplementary Note 1.1) and was therefore not included in further analyses. While both monkeys typically observed several shapes before making their decisions, monkey S used more shapes than monkey T (Fig. 2b, mean number of used shapes: 7.47 for monkey S, 2.75 for monkey T). Hence, monkey S gathered more information about the correct target, leading to higher response accuracy (fraction correct choice: 0.90 for monkey S, 0.65 for monkey T). Thus, while observing more shapes can, in principle, lead to higher accuracy, the two monkeys used different strategies, with monkey T opting for fewer observations at the cost of reduced decision accuracy.

**Figure 2.**
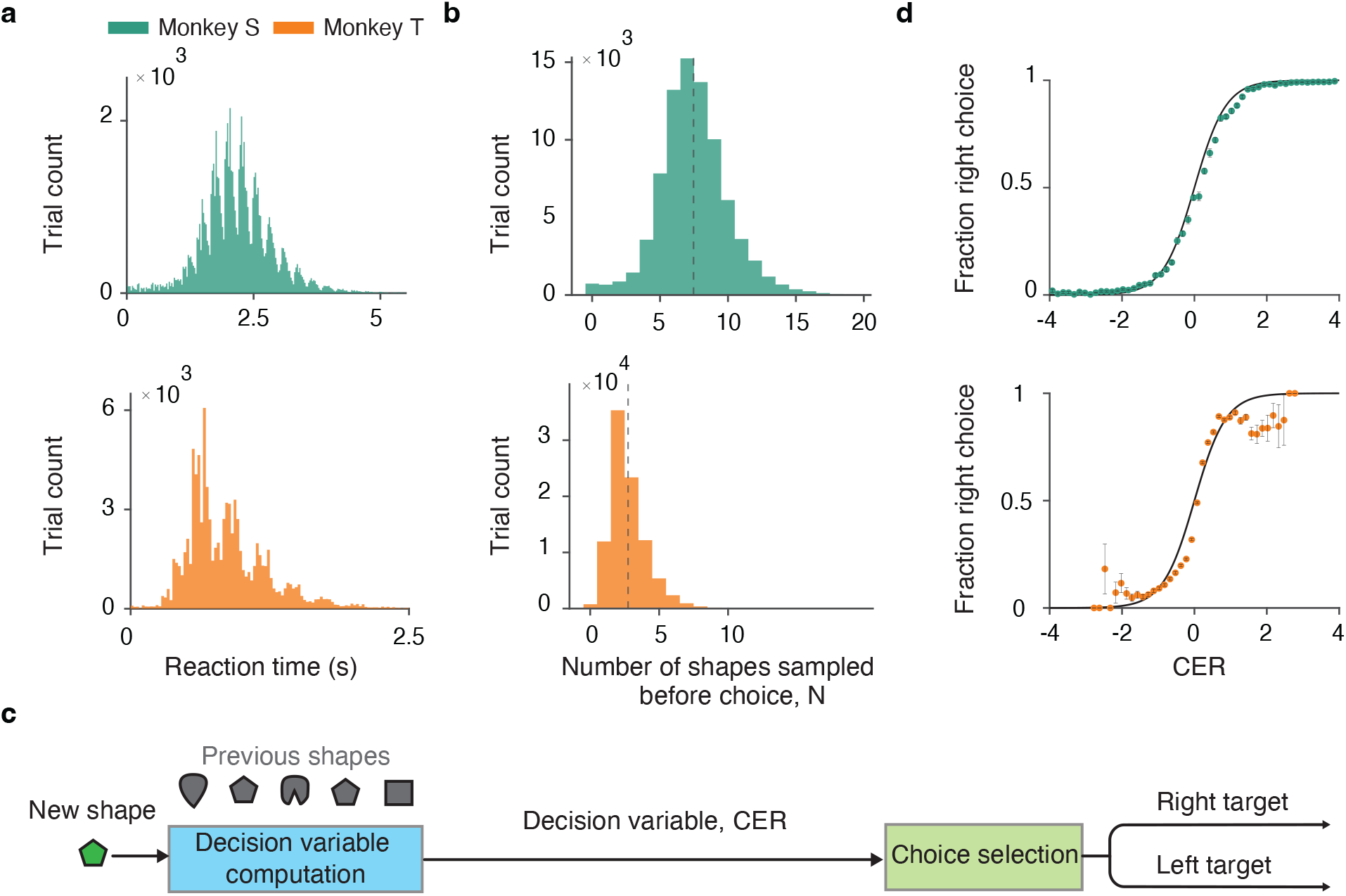
Behavioral performance in the sequential probabilistic reasoning task. **a**, Reaction time histogram (bin size 25 ms) for monkey S (green) and monkey T (orange). **b**, Histogram of the number of shapes *N* used by each monkey. For each trial, *N* does not include the shape displayed during the saccade. Dashed lines indicate mean values. **c**, The sequential probabilistic reasoning task involves two computational components: computing a decision variable (blue box) and selecting a choice based on this decision variable (mint box). **d**, The probability matching strategy sets the choice probability equal to the correct Bayesian posterior probability of the right choice given the observed cumulative evidence (black line). Dots indicate the proportion of trials in which the monkey chose the right target at each cumulative evidence level, with cumulative evidence binned in 0.15 intervals. Error bars represent standard deviation in each bin.

We modeled the monkeys’ choices to uncover behavioral strategies underlying their differing decision accuracy. In the probabilistic reasoning task, the decision-making process involves two components: (1) computation of a decision variable from the sequence of observed shapes, and (2) a choice selection strategy generating categorical choices based on the value of the decision variable (Fig. 2c). We inferred these two decision components separately. First, we determined the monkeys’ choice selection strategy, assuming the decision variable of an ideal Bayesian observer. With the choice selection strategy established, we then refined our quantification of the decision variable the monkeys used to guide their choices.

### Choice selection strategy

To infer the monkeys’ choice selection strategy, we assumed that the decision variable represents the true cumulative evidence from all observed shapes, reflecting perfect evidence accumulation as in the ideal Bayesian observer. Since the shapes are sampled independently, their cumulative evidence for the right target (CER) is the sum of their true weights 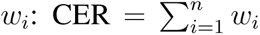. Thus, after observing *n* shapes, the probability of the right target being rewarded is a logistic function of the cumulative evidence (Fig. 2d, Supplementary Note 1.2):

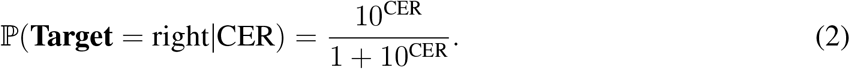

The choice selection strategy defines how the reward probability maps onto the choice. An ideal Bayesian observer always chooses the target that has higher reward probability, that is, choosing right if ℙ(**Target** = right|CER) *>* 0.5 and otherwise choosing left. When the cumulative evidence is contaminated by noise, this strategy results in behavior mimicking probability matching^42^, in which the monkey’s choice probabilities align with the anticipated reward probabilities. Probability matching is commonly observed in both humans^43, 44^ and rhesus monkeys^45, 46^. The behavior of our monkeys also approximately followed probability matching (Fig. 2d), indicating that the animals learned and adequately used the weights of the shapes.

However, the monkeys’ behavior slightly deviated from probability matching, which was especially noticeable at the extreme values of the cumulative evidence for monkey T (Fig. 2d). We hypothesized that these deviations, though seemingly small, reflect working memory limitations that lead to suboptimal estimation of the decision variable, such that the monkey’s subjective cumulative evidence differs from that of an ideal Bayesian observer. To test this hypothesis, we developed behavioral models that incorporate working memory limitations in the computation of the decision variable.

### Suboptimal evidence accumulation

We investigated the sources of deviations from optimal evidence accumulation in monkey behavior. First, we tested whether the monkeys’ learned subjective shape weights *w̃_i_* differed from the true weights *w_i_*. To estimate the subjective weights *w̃_i_*, we fitted monkeys’ behavior with a model in which the choice probability depends on the subjective cumulative evidence toward the right target 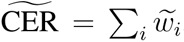, where the subjective weights of individual shapes are free parameters (Methods). All shapes had nonzero inferred subjective weights (Fig. 3a), indicating that the monkeys incorporated every shape into their decisions rather than ignoring any specific shape. For the eight non-trump shapes, the inferred subjective weights tightly corresponded with the true weights (monkey S: *R*^2^ = 0.94, *p* = 10^−4^; monkey T: *R*^2^ = 0.92, *p* = 2 · 10^−4^). The subjective weights of the trump shapes had the correct signs but substantially lower magnitude than their true weights, consistent with previous findings^38, 39^. This under-weighting likely results from low frequency of trump shapes, which makes their precise values more difficult to learn. Since both monkeys accurately learned the weights of all non-trump shapes, other factors likely contributed to the observed deviations from optimal evidence accumulation.

**Figure 3.**
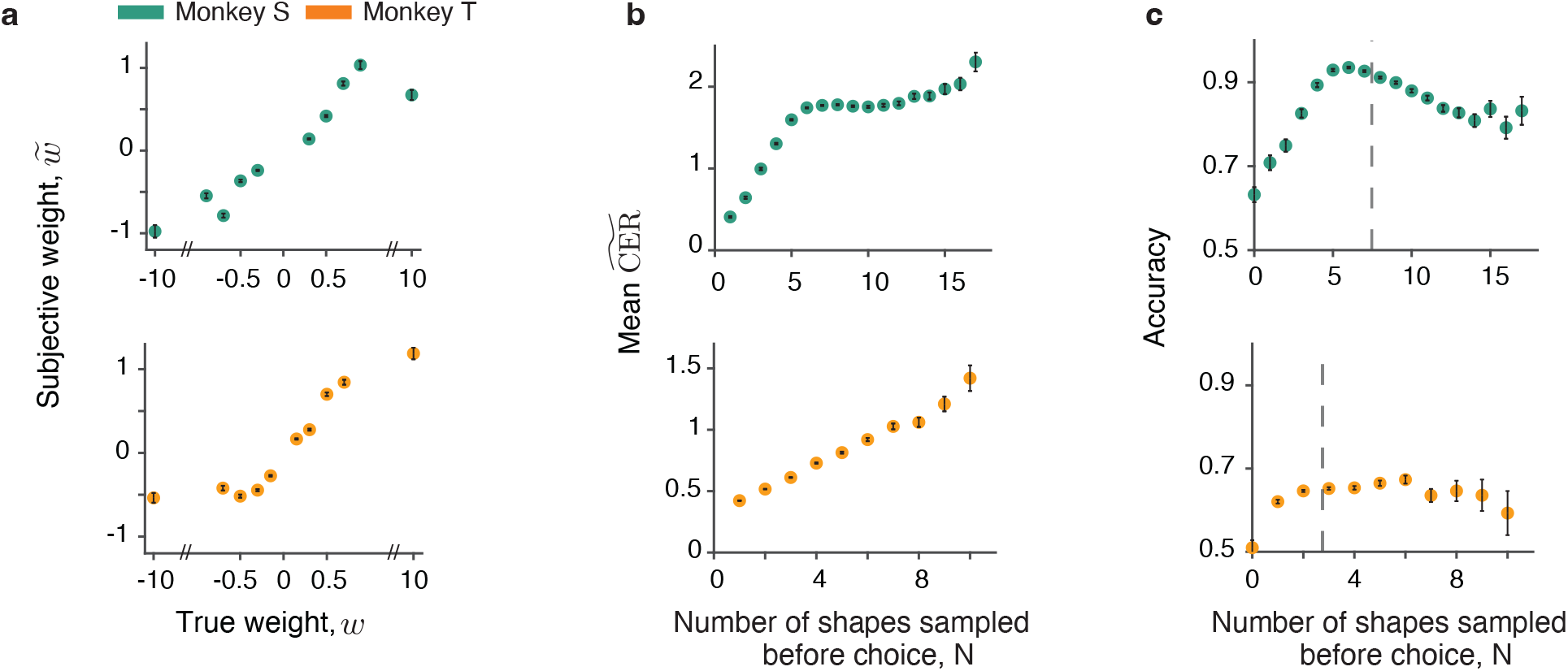
Behavior deviates from perfect evidence accumulation. **a**, The subjective weights estimated under the assumption of perfect evidence accumulation (y-axis) closely correspond to the true shape weights (x-axis). Error bars represent 95% confidence interval. **b**, Average cumulative evidence at decision time as a function of the number of shapes used to make a choice. Error bars represent standard error of the mean (SEM). **c**, Decision accuracy as a function of the number of shapes used to make a choice. Error bars represent standard error of the mean (SEM). Accuracy initially improved as more shapes were sampled, then plateaued and eventually declined in both monkeys. The decline in accuracy after many shapes had been sampled indicates that the monkeys did not use all available evidence and therefore did not accumulate evidence perfectly.

We therefore tested whether the monkeys used all available evidence to inform their choices, as expected under perfect evidence accumulation. If the monkeys perfectly integrated subjective weights of all shapes, the subjective cumulative evidence 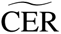 at decision time would correlate with the decision accuracy: choices should be more accurate when more evidence is available. Our data were inconsistent with this scenario. 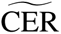 at decision time increased with the number of sampled shapes, growing non-linearly for monkey S and more linearly for monkey T (Fig. 3b). In contrast, the decision accuracy initially rose, then plateaued, and eventually declined for both monkeys as the number of sampled shapes *N* increased (Fig. 3c). This inconsistency between the increasing 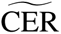 and decreasing accuracy at large *N* (above ∼ 6 for monkey S and ∼ 3 for monkey T) indicates that the monkeys did not use all available evidence to inform their choice, suggesting that part of this evidence was effectively lost. Thus, for large *N*, the monkeys’ subjective 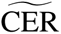 deviated from the perfect evidence accumulation.

### Working memory limitations constrain decision-making

The monkeys’ failure to use all available evidence led us to hypothesize that their decision-making was affected by working memory limitations. To test this hypothesis, we fitted animals’ choices with a model that incorporated working memory limitations (WML), including a constant-rate memory decay, serial-position biases—primacy^8, 9^, or differential weighting of the first item, and recency^10, 11^, or differential weighting of the last item—and a choice-history priming^12, 13^, a bias toward repeating the previous choice. In the WML model, the subjective weights of sampled shapes are scaled by exponential temporal decay factors and by primacy and recency factors determined by each shapes’ position in the sequence. The scaled weights are then summed with an added choice-history bias to compute the subjective cumulative evidence CER_WML_ (Methods). The WML model explained monkey’s choices significantly better than the perfect evidence accumulation model (Fig. 4a, difference in Akaike information criterion, AIC: 851 for monkey S; 2,577 for monkey T), supporting our hypothesis that suboptimal decision-making resulted form working memory limitations.

**Figure 4.**
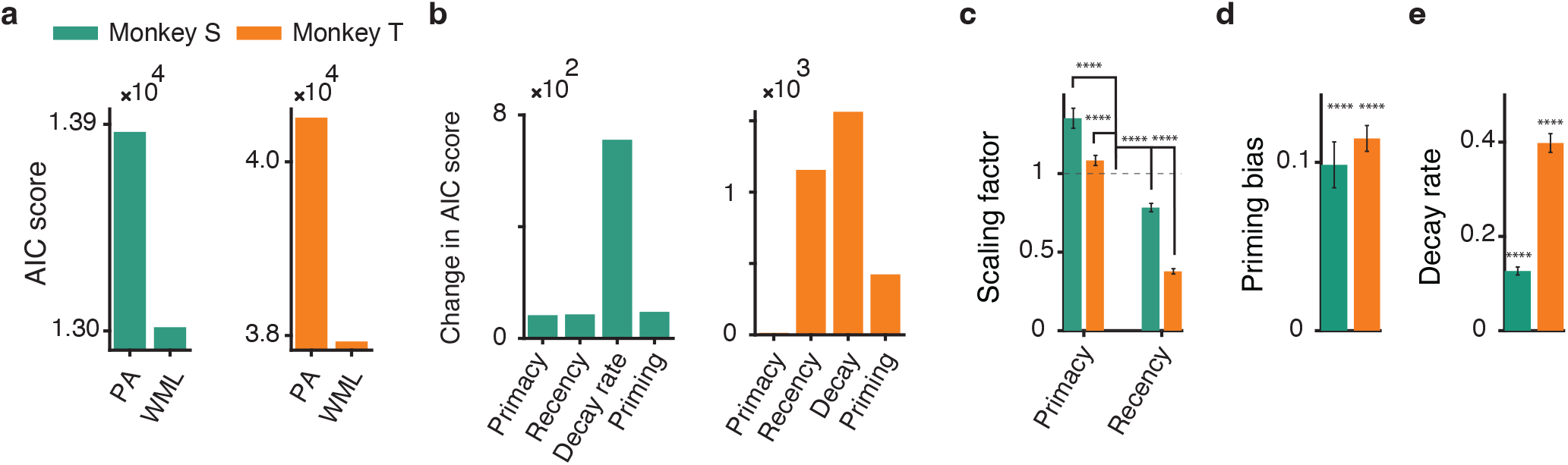
Working memory limitations account for suboptimal behavior. **a**, The WML model accounts for the behavior better than perfect evidence accumulation (PA), as indicated by lower AIC scores for both monkey S (green, PA 13,867; WML 13,016) and monkey T (orange, PA 40,508; WML 37,931). **b**, Each working memory constraint (memory decay, primacy, recency, and priming) contributes significantly to behavior, as indicated by the increase in AIC scores when each constraint is removed individually from the WML model. Memory decay has the greatest impact, producing the largest increase in AIC score when removed. **c**, The first and last shapes in the sequence contribute to cumulative evidence with their subjective weights multiplied by the primacy and recency scaling factors, respectively, whereas all intervening shapes are assigned the baseline factor of 1. In both monkeys, the primacy factor was significantly greater than 1, whereas the recency factor was significantly less than 1 (^∗∗∗∗^*p <* 10^−10^). **d**, Both monkeys exhibit a significant positive priming bias, indicating the tendency to repeat their previous choice, regardless of the prior trial outcome (^∗∗∗∗^*p <* 10^−10^). **e**, Both monkeys exhibit significant memory decay, with the decay rate approximately threefold higher in monkey T than monkey S (^∗∗∗∗^*p <* 10^−10^).

To determine the unique contribution of each factor, we systematically removed individual components from the WML model and assessed the resulting change in AIC scores. All four working memory limitations—priming, primacy, recency, and decay—significantly improved the model performance, with memory decay having the largest impact in both monkeys (Fig. 4b). Specifically, the first observed shape contributed significantly more to subjective cumulative evidence than subsequent shapes, whereas the last shape contributed significantly less (Fig. 4c). Additionally, both monkeys displayed a bias toward repeating their previous choice, indicating a priming effect (Fig. 4d).

The difference in memory decay rates between the two monkeys explained the difference in the number of shapes they sampled before reaching a decision. The decay rate *α* was threefold higher in monkey T than in monkey S (Fig. 4e). This disparity in decay rates can explain why monkey T used fewer shapes on average than monkey S (Fig. 2b, monkey T 2.75, monkey S 7.47). Theoretically, in the absence of memory decay, the average incremental evidence gain Δ(*n*) after observing the *n*th shape remains constant and independent of 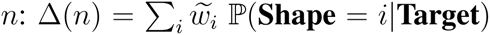 which implies that each additional shape provides the same expected increase in cumulative evidence. However, with decay, Δ(*n*) ∝ exp(−*αn*) (Supplementary Note 1.3), such that the expected gain in cumulative evidence from an additional shape decreases as more shapes are observed. If waiting carries a cost, the optimal decision policy is to terminate evidence accumulation when the marginal benefit of sampling an additional shape falls below that cost. Our results support this trade-off: monkey T, with a higher decay rate, terminated the decision process earlier, while monkey S, with a lower decay rate, sampled more shapes to maximize information and improve accuracy.

The WML model accounts for critical behavioral patterns that the perfect evidence accumulation cannot explain. First, the perfect evidence accumulation fails to explain why accuracy does not consistently increase with the number of observed shapes (Fig. 3c). Second, for monkey T, choices deviate substantially from probability matching at extreme values of cumulative evidence when the decision variable is computed under the assumption of perfect evidence accumulation (Fig. 2d). These deviations occur because the perfect accumulation overestimates subjective cumulative evidence at extreme values. By incorporating working memory limitations, the WML model brings the choices of both monkeys into close agreement with probability matching (Fig. 5a). Third, the distinct, non-monotonic dependence of accuracy on the number of observed shapes (Fig. 3c) closely mirrors the non-monotonic dependence of the subjective cumulative evidence at decision time in both monkeys (Fig. 5b), restoring the expected relationship between cumulative evidence and decision accuracy. Together, these findings demonstrate that the monkeys’ deviations from optimal decision-making arise from working memory limitations that affect how evidence is retained, weighted, and accumulated over time.

**Figure 5.**
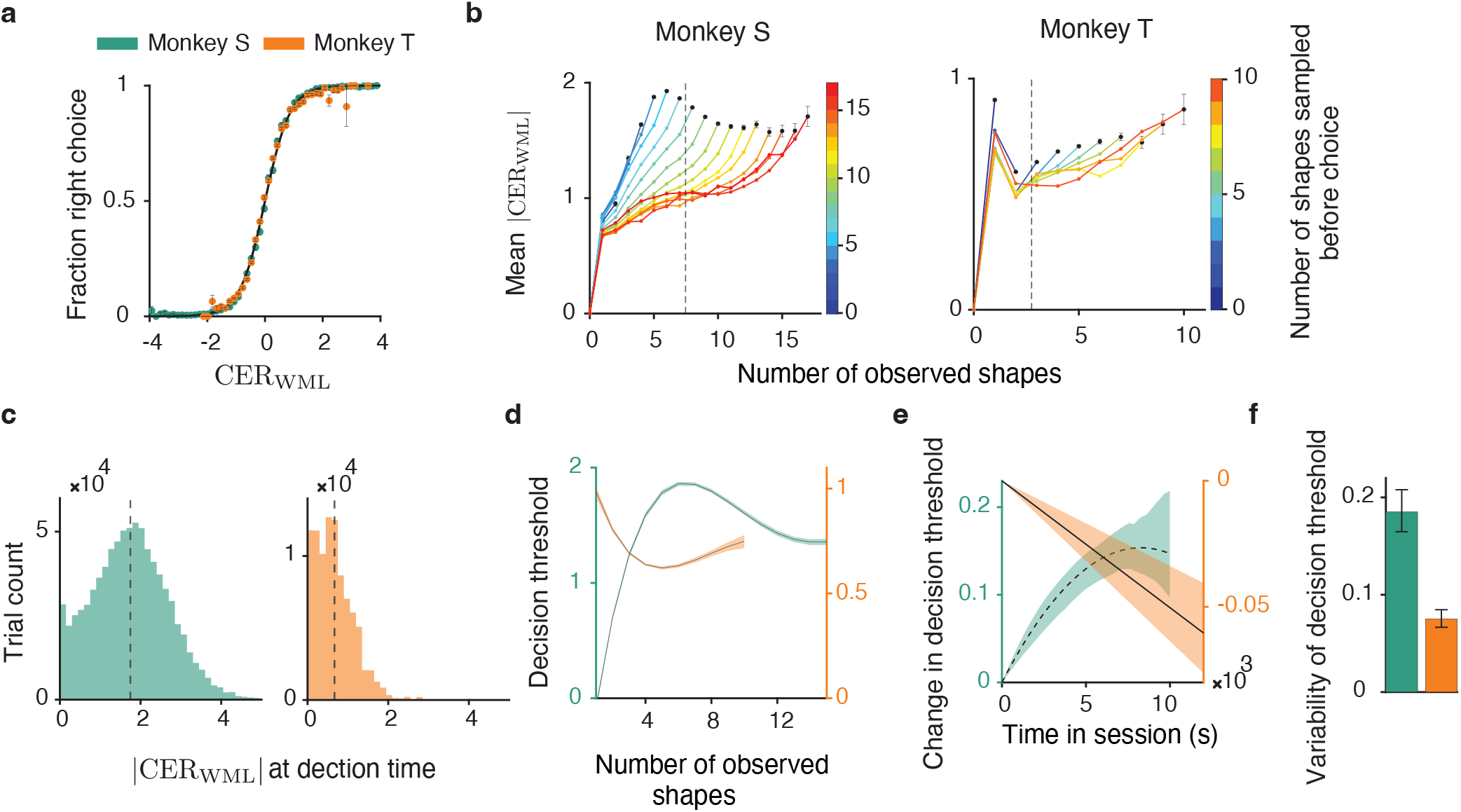
Stopping rule for evidence accumulation. **a**, The Bayesian posterior probability of the right choice given subjective cumulative evidence from the WML model (black line) closely matches the probability that the monkey chose the right target at each evidence level (dots; subjective cumulative evidence binned in 0.15 intervals). This agreement shows that, after accounting for working memory limitations, the monkeys follow probability matching strategy. Error bars indicate the standard deviation within each bin. **b**, Average absolute value of subjective cumulative evidence after each newly observed shape, plotted separately for trials with different total numbers of shapes sampled before choice, indicated by color, for monkey S (left) and monkey T (right). **c**, Histogram of the absolute value of subjective cumulative evidence at decision time from the WML model for monkey S (green) and monkey T (orange). Dashed lines indicate the mean values. **d**, Decision threshold as a function of the total number of shapes sampled before choice, estimated using a nonlinear mixed-effects model for monkey S (green) and monkey T (orange). The shading indicates 95% confidence interval. **e**, Change in decision threshold as a function of time in session, estimated using a nonlinear mixed-effects model for monkey S (green) and monkey T (orange). The shading indicates 95% confidence interval. **f**, Standard deviation of session-by-session variability of the decision threshold, estimated using a nonlinear mixed-effects model for monkey S (green) and monkey T (orange). The error bars indicate 95% confidence interval.

### Stopping rule for evidence accumulation

As the WML model accurately captured the monkeys’ behavior, it allowed us to examine the stopping rule for evidence accumulation, that is, how the animals determined when sufficient evidence had been collected to commit to a choice. An optimal Wald strategy prescribes accumulating evidence until it reaches a fixed threshold, which may be specific to each individual^39, 47^. This strategy achieves a desired accuracy, set by the decision threshold, using the minimum expected number of samples. Under Wald’s strategy, the absolute subjective cumulative evidence |CER_WML_| at decision time should remain nearly constant, regardless of the number of observed shapes. However, inconsistent with this prediction, the average |CER_WML_| showed systematic dependence on the number of observed shapes in both animals (Fig. 5b). Monkeys S showed an initial ‘optimism’ bias: it committed to a choice immediately when the first few shapes produced relatively high |CER_WML_|, reflected in lower |CER_WML_| value at decision time. For monkey S, sampling more than six shapes triggered an urgency effect, reflected in a collapsing decision threshold, whereas the threshold increased with the number of sampled shapes for monkey T. Thus, the monkeys’ stopping rule deviated from the fixed-threshold policy prescribed by Wald’s strategy.

The deviation from the Wald’s strategy was also evident in the distribution of |CER|_WML_ at decision time, which quantifies how frequently the monkeys committed to a choice at different levels of accumulated evidence (Fig. 5c). Under Wald’s strategy, this distribution should be narrow and peaked just above the evidence threshold, with variability arising only from the overshoot in accumulated weights required to cross the fixed threshold. This variability is bounded by the standard deviation of the absolute subjective weights |*w̃_i_*| across all shapes (*σ* = 0.19 for monkey S and *σ* = 0.16 for monkey T). However, the observed standard deviation of |CER| at decision time was substantially larger (0.92 for monkey S, 0.47 for monkey T), further reinforcing the deviation from Wald’s strategy.

To disentangle factors that contribute to the observed variability in the decision threshold, we used a non-linear mixed-effects model (Methods). The model fitted the dependence of |CER| at decision time on the number of observed shapes (e.g., due to urgency as the time cost of gathering information increases^48^), time within a session (e.g., due to shifts in impulsivity), and random variation across sessions. For both monkeys, the decision threshold varied with the number of observed shapes (Fig. 5d), consistent with the dependence observed in the average |CER_WML_| (Fig. 5b). This analysis confirmed that monkey S exhibited initial optimism when the first few shapes produced strong evidence, reflected by a lower decision threshold, with urgency emerging at larger number of observed shapes, as the decision threshold decreased. Monkey T, on the other hand, displayed urgency immediately after the first shape, and later the threshold increased with the number of sampled shapes. The decision threshold also changed over the course of a session, rising for monkey S and decreasing for monkey T (Fig. 5e). Both monkeys also exhibited significant session-to-session variability in the decision threshold (Fig. 5f). Together, these findings reveal that monkeys’ decisions were affected by both working memory distortions in the accumulated evidence and a dynamic stopping threshold that determined how much evidence was required to commit to a choice.

### A circuit mechanism of memory decay

We developed a neural circuit model to investigate possible mechanisms underlying the memory decay that affected evidence accumulation observed in the monkeys’ decision-making behavior. At single-neuron level, the decay timescale is influenced by membrane and synaptic time constants. For a typical cortical neuron, the membrane time constant is approximately 5 − 20 ms^49, 50^. In the absence of synaptic input, the membrane potential decays to its resting potential within this brief window. Similarly, synaptic time constants for excitatory cortical neurons typically range between −150 ms^49, 51^. However, the working memory decay timescale for both monkeys was significantly longer, on the order of seconds, suggesting that the slower memory decay observed in behavior arises from the collective dynamics of interacting neurons, rather than being governed solely by the intrinsic properties of individual neurons.

We trained a neural circuit model to reproduce the choice behavior of each monkey (Fig. 6a). The model consisted of two units integrating evidence for the right and left choices. The difference in their firing rates defined the cumulative evidence, which determined choice probability through a logistic function Eq. 2. Each unit incorporated a leak term and self-excitation with strength *s*_exc_. The unit’s effective time constant is given by *τ/*(1 − *s*_exc_), extending beyond its intrinsic time constant *τ* (Methods). Thus, by adjusting *s*_exc_, self-excitation allows the model to capture a range of behavioral timescales. When *s*_exc_ = 1, the circuit functions as a perfect integrator. As self-excitation decreases, the memory decays more rapidly. The integrator units received inputs from a layer of ten shape-selective inputs, each activated by one shape^52^. The connection weights from the input to integrator units and the self-excitation parameter were optimized to match the monkey behavior (Methods).

**Figure 6.**
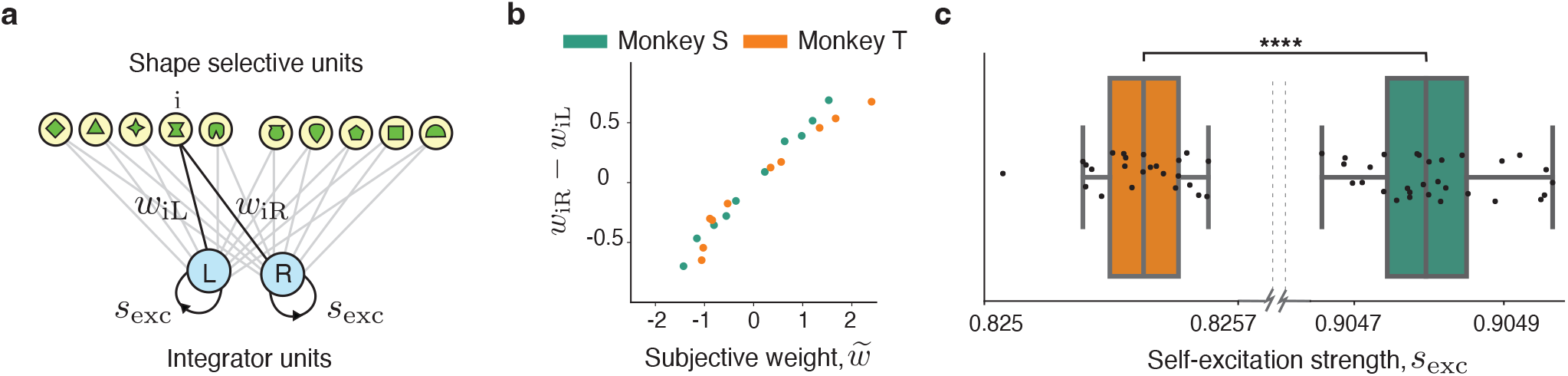
A circuit model of memory decay. **a**, The model consists of two units integrating evidence for the right (R) and left choices (L). These units incorporate self-excitation with strength *s*_exc_. The integrator units receive inputs from a layer of ten shape-selective units, each activated by one shape. The connection weights from the input to integrator units (*w_iR_*and *w_iL_*) and self-excitation parameter *s*_exc_ are optimized to match the monkey behavior. **b**, The subjective shape weights (*w̃_i_*, x-axis) are linearly related to the difference in connection weights from the corresponding input to the left and right integrator units (*w_iR_* − *w_iL_*, y-axis), averaged over initializations of the trained networks. **c**, In trained networks, self-excitation parameter *s*_exc_ is significantly larger for monkey T (green) than for monkey S (orange), explaining the differences in their memory decay rate. Dots show 30 initializations of the network. In boxplots, the center lines indicate medians, the boxes span the 25–75th percentiles, the whiskers extend to the nearest of 1.5× the inter-quartile range (^∗∗∗∗^*p <* 10^−10^).

The circuit model fitted the behavior better than the perfect evidence accumulation model (AIC score, monkey S: 13,486; monkey T: 38,887), but less accurately than the WML model (Fig. 4a), because the circuit model did not implement the primacy, recency, and priming mechanisms included in the WML model. The connection weights from the shape-selective inputs to the integrator units closely matched the subjective weights in the WML model (Fig. 6b), suggesting that subjective weights can be learned through synaptic plasticity at the inputs to the evidence-accumulation circuit^52^. The fitted self-excitation parameter was less than one in both animals (Fig.6c; monkey S: 0.9048 ± 0.0009, monkey T: 0.8254 ± 0.0009; mean ± std across initializations), accounting for the memory decay. The self-excitation was larger in monkey S than in monkey, consistent with the faster memory decay in monkey T. These results show that the observed working memory decay is consistent with a circuit mechanism in which self-excitation facilitates evidence integration over extended timescales, while learned inputs weights determine the relevant weighting of sensory cues.

### Dopaminergic modulation of working memory and evidence integration

To determine how dopaminergic signaling modulates working memory constraints during sequential decision-making, we examined the effects of pharmacological interventions on the monkeys’ behavior. Dopamine plays critical computational roles in decision-making that relies on working memory by selectively gating sensory input to prioritize relevant information^53^, maintaining and manipulating working memory contents^22, 36^, and modulating motor commands to translate decisions into actions^54^. At the circuit level, optimal dopamine levels stabilize neural representations in PFC, supporting the maintenance and manipulation of information by enhancing synaptic connectivity and neuronal responsiveness through D1 and D2 receptor activity. We systemically administered either SCH-23390 (a D1R antagonist), SKF-81297 (a D1R agonist), or saline (control) mid-session to investigate how dopamine modulates the various components of sequential decision-making, including subjective weights, working memory decay, recency, primacy, and priming bias.

Following the injection of a D1R antagonist, both monkeys integrated significantly more shapes into their decisions compared to the control condition (Fig. 7a, Methods). At the high dose, monkeys S and T sampled an average of 1 and 0.5 additional shapes, respectively. Despite this extended shape sampling, the performance of both monkeys remained largely unchanged (monkey S: no drug 0.901 ± 0.001, saline 0.909 ± 0.003, SCH 0.909 ± 0.003, SKF 0.906 ± 0.004; monkey T: no drug 0.646 ± 0.002, saline 0.661 ± 0.005, SCH 0.667 ± 0.006, SKF: 0.660 ± 0.005). Thus, while D1R blockade increased the amount of evidence sampled, it did not translate to improved task performance.

**Figure 7.**
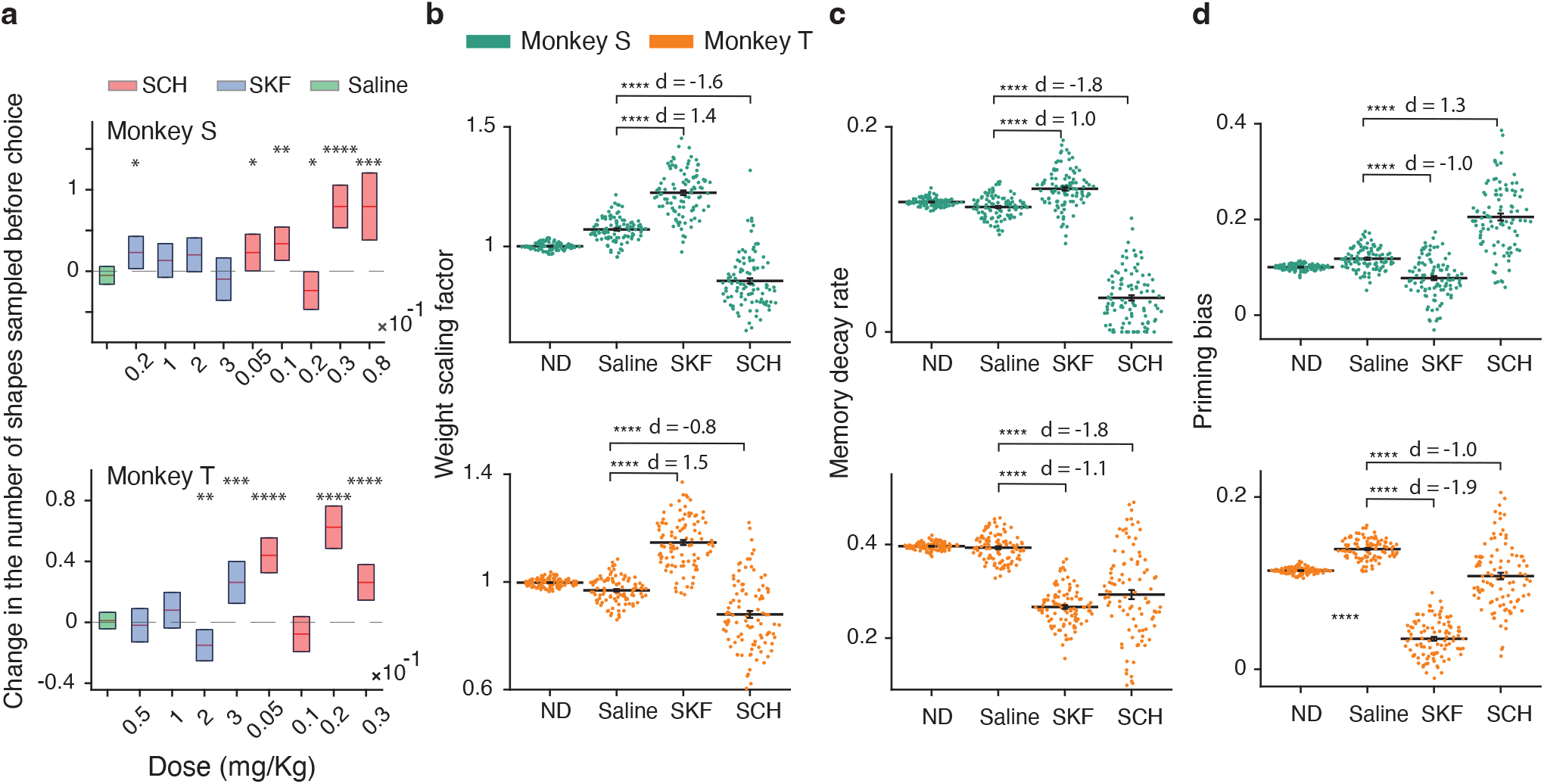
Dopamine modulation of working memory constraints. **a**, Change in the number of shapes sampled before choice under saline (Saline), dopamine agonist (SKF), and dopamine antagonist (SCH) conditions at varying doses, relative to no-drug (ND) condition for monkey S (upper panel) and monkey T (lower panel). The number of trials in each condition was: ND 81,985; Saline 7,287; SCH 6,818; SKF 6,152. Both monkeys used significantly more shapes under the dopamine antagonist injection, whereas saline and the dopamine agonist injections did not produce a consistent effect across doses and animals. The box indicates 95% confidence interval, the center line indicates the median across sessions. The p-values are from two-tailed t-test (^∗^*p <* 0.05, ^∗∗^*p <* 0.01, ^∗∗∗^*p <* 10^−3^, ^∗∗∗∗^*p <* 10^−10^). **b**, Effects of drug injections on subjective evidence weighting. The weight-scale factor is a multiplicative factor by which drug injection scales all subjective weights, with a value of 1 indicating no drug effect. In both monkeys, dopamine agonist injection significantly increased the subjective weights relative to saline, with large Cohen effect sizes (|*d*| *>* 1.3). Conversely, in both monkeys, dopamine antagonist injection significantly reduced the subjective weights relative to saline, with large Cohen effect sizes (|*d*| *>* 0.8). The data were pooled across SKF dosages ⩾ 0.2 and SCH dosages ⩾ 0.03 (^∗∗∗∗^*p <* 10^−10^). **c**, The dopamine antagonist significantly reduced memory decay rate in both monkeys, with very large Cohen effect sizes (|*d*| *>* 1.3). The effects of the dopamine agonist varied by monkey: monkey T exhibited a significant reduction in the decay rate, whereas monkey S showed no significant effect. Saline had no effect for either monkey (^∗∗∗∗^*p <* 10^−10^). **d**, The dopamine agonist significantly reduced the priming bias in both monkeys, with large Cohen effect sizes (|*d*| *>* 0.8). The effects of the dopamine antagonist varied by monkey: monkey S showed a significant increase in the priming bias, whereas monkey T showed no significant effect (^∗∗∗∗^*p <* 10^−10^). In a–c, each point represents a model fit for one bootstrap sample, with 100 bootstrap samples used in each condition.

Increased sampling could be driven by reduced subjective evidence weighting, but additional factors may also contribute. To disentangle these factors, we used the WML model and fitted the five free parameters—primacy, recency, decay, priming, and a weight-scale factor—in control condition and when monkeys were affected by DA challenges. To limit the number of free parameters, we assumed the subjective weights of the shapes retained the same relative magnitudes as those estimated in the no-drug condition, but were scaled by a global weight-scale factor, with a value of one indicating no drug effect. For improved statistical power, the data from SKF doses ⩾ 0.2 and SCH doses ⩾ 0.03 were combined.

The D1R agonist significantly increased the weight-scale for both monkeys, with large effect sizes (Fig. 7b, monkey S, *p <* 10^−10^; monkey T, *p* =*<* 10^−10^; Cohen’s size effect: *d >* 1.3, 100 bootstrap samples, two-tailed t-test). Conversely, the D1R antagonist significantly reduced the weight-scale, also with large effect sizes (Fig. 7b, monkey S: *p <* 10^−10^; monkey T: *p* = 6 × 10^−10^; Cohen’s size effect: *d >* 1.3, 100 bootstrap samples, two-tailed t-test). These findings show that dopamine bidirectionally modulates the subjective evidence weighting: D1R agonists increased the perceived importance of shapes, whereas and D1R antagonists reduced it.

D1R manipulation further influenced working memory stability by altering the rate of memory decay. The D1R antagonist reduced working memory decay in both monkeys (Fig. 7c; monkey S, *p <* 10^−10^; monkey T, *p <* 10^−10^; large Cohen’s effect size: |*d*| *>* 0.8, 100 bootstrap samples, two-tailed t-test), consistent with both animals sampling more shapes under the D1R antagonist (Fig. 7a), when additional evidence can be retained longer. While this result might seem counterintuitive, given that dopamine typically facilitates working memory by stabilizing information in PFC^36, 55, 56^, it is consistent with the ‘inverted-U’ hypothesis, where both insufficient and excessive dopamine levels impair working memory performance^28, 57^. This interpretation is supported by the observation that the effect of the D1R agonist on working memory decay differed between the two monkeys (Fig. 7c), suggesting they may have occupied different baseline positions on the dopamine-response curve. Specifically, D1R activation reduced memory decay in monkey T, which exhibited larger baseline decay, but increased decay in monkey S, which exhibited smaller baseline decay, consistent with the idea that the same dopamine manipulation can improve or impair memory stability depending on baseline dopamine state. Beyond memory decay, the D1R agonist also decreased the bias toward repeating the last choice for both monkeys (Fig. 7d, monkey S, *p <* 10^−10^; monkey T, *p <* 10^−10^; large Cohen’s effect size: |*d*| *>* 0.8, 100 bootstrap samples, two-tailed t-test). Thus, prior choices had less influence on current decisions following D1R activation. DA manipulations also affected the primacy and recency factors (Supplementary Note 1.4).

In summary, our results demonstrate that D1R signaling plays a pivotal role in modulating the computational components of sequential probabilistic reasoning. D1R blockade simultaneously reduced working memory decay rates and lowered subjective weights, while D1R activation increased subjective weights and attenuated the priming bias from previous choices. Collectively, these findings suggest that dopamine regulates sequential probabilistic reasoning by fine-tuning working memory limitations, specifically by shifting the balance between memory decay, evidence valuation, and the influence of prior choices.

## Discussion

Working memory decay, primacy, recency, and priming all constrain decision-making when probabilistic evidence needs to be integrated over extended time. working memory decay further affects individual stopping-rule strategies for evidence accumulation, as forgetting offsets the evidence gained from sampling additional shapes. Dopamine modulates these constraints by altering the gain of sensory evidence, the rate of memory decay, and the bias towards repeating prior choices.

Our models reveal that animals’ behavior in the probabilistic reasoning task deviates substantially from the perfect evidence accumulation with an optimal fixed-threshold stopping rule. A previous study using a nearly identical probabilistic reasoning task^39^ found that, on average, cumulative evidence at decision time was similar whether monkeys based their choice on few or many shapes, reminiscent of a fixed-threshold Wald strategy. However, cumulative evidence at decision time varied substantially across trials, with the standard deviation reaching three to four times the mean, a pattern interpreted as possibly reflecting noise^39^. Our work suggests an alternative explanation: deviations from optimality resulting from working memory constraints. Accounting for these constraints restores the close relationship between cumulative evidence and decision accuracy and aligns animals choices with probability matching. Although previous work did not consider working memory constraints, it also reported substantial individual differences between monkeys, with one animal routinely sampling many shapes and the other sampling relatively few^39^. Our results suggest that these individual differences may also have arisen from differences in memory-decay rates between animals.

Although the monkeys’ behavior deviated from the ideal Wald strategy, it was nearly optimal given their working memory limitations. If information from more than *n* shapes ago is irretrievably lost, sampling beyond *n* shapes provides little behavioral benefit. Sampling behavior of both monkeys was consistent with their rates of memory decay, with the animal showing slower decay sampling for longer. Additionally, one monkey exhibited early optimism, committing to a choice quickly if initial evidence was strong, while both monkeys exhibited urgency, reflected in a decreasing decision threshold after several shapes had been sampled. Previous studies have proposed possible neural circuit mechanisms underlying early optimism and urgency^58–61^, but further work is needed to integrate working memory limitations into such circuits. Our neural circuit model suggests that the strengths of self-excitation within decision-making circuits determines the rate of memory decay, consistent with previous modeling and experimental work^16, 21, 35^. Recency and primacy effects have been attributed to attentional processes^62^. Priming, that is, the tendency to repeat the previous choice, is common behavior even when it does not benefit performance^63–66^. Priming effects may arise from residual activity or short-term synaptic plasticity in decision-related neural populations carrying the previous choice or decision variable, which can bias subsequent evidence accumulation^67–69^, or from biases in the evidence-accumulation process itself^70^.

We observed that dopamine modulates subjective weights in both monkeys. Given their extensive training, it is likely that these weights were encoded in long-term memory. One possibility is that dopamine adjusts the encoding of long-term memory, a mechanism previously observed in rats^71^. Alternatively, dopamine may not alter the weights themselves but instead modulate synaptic efficacy, thereby affecting the retrieval of weights from long-term memory. This mechanism has been demonstrated in studies with mice and humans^72, 73^. Further neural recordings are necessary to distinguish between these possibilities.

The effect of dopaminergic manipulation on decision-related weighting may have relevance for clinical conditions. Recent studies have shown that patients with obsessive-compulsive disorder (OCD) tend to underestimate subjective weights in a sequential probabilistic reasoning task similar to ours^47^. As a result, OCD patients are less effective at transforming sensory information into evidence. However, they maintain similar accuracy to healthy controls by taking longer to respond, i.e., by sampling more evidence. This behavior resembles the monkeys’ performance under dopamine antagonists. The aetiology of OCD is not fully established, although cortico-striatal-thalamo-cortical (CSTC) circuit dysfunction has been proposed as one possible contributor to symptoms^74–77^. The underlying OCD neurobiology is complex, with serotonin, dopamine and glutamate dysregulation all implicated as potentially relevant^78^. Selective serotonin reuptake inhibitors are partially effective as a treatment, sometimes in conjunction with dopamine-receptor-blocking antipsychotics, suggesting involvement of both serotonergic and dopaminergic mechanisms. Our findings are consistent with the possibility that dopaminergic dysfunction may contribute to altered evidence weighting in OCD. In our task, D1R blockade resulted in reduced subjective weighting, unexpectedly reduced memory decay, and an increased tendency to repeat the previous choice. By contrast, increased D1R activation produced the opposite pattern. Our models provide a reliable analytical framework for dissociating how neuromodulation affects distinct components of decision-making in health and psychiatric disease.

In summary, our work advances the understanding of decision-making processes that rely on working memory and identifies mechanisms that may be relevant to clinical interventions in conditions like OCD and other cognitive disorders. Future research, particularly work examining the neural circuit underpinnings of these limitations across brain areas, will be essential for elucidating the mechanisms behind working memory limitations and integrating them into broader frameworks of cognitive function.

## Methods

### Procedures and animals

All surgical and behavioral procedures conformed to the UK Animals Scientific Procedures Act, European Communities Council Directive RL 2010/63/EC, and the U.S. National Institutes of Health Guidelines for the Care and Use of Animals for Experimental Procedures.

One male and one female adult rhesus monkey (8.9 and 11kg) were trained to comfortably sit in a plexiglass chair. Before training, they had been implanted with a head holder device under sterile conditions. Details of surgical procedures, postsurgical management, and analgesics have been described previously^79^. To promote behavioral motivation and performance, daily access to fluid was controlled during training and experimental periods, using fluid control regimes with minimal psychological impact and no measurable physiological impact on the animals^80^.

Stimulus presentation and behavioral task control were achieved through Remote Cortex 5.95 (Laboratory of Neuropsychology, National Institute for Mental Health, Bethesda, MD). Task stimuli were presented on a cathode ray tube (CRT) monitor (120Hz, 1280 × 1024 pixels). Raw behavioral data were collected through Remote Cortex 5.95. Eye movements and pupil diameter were tracked and recorded using an SR Research EyeLink eye tracker (Eye-Link version 2.04, SR Research/SMI). For all experimental sessions, animals performed the task in a dark room sitting upright in primate chairs.

### Probabilistic reasoning task

Animals performed the Sequential Probability Ratio Task^39^, referred to as the probabilistic reasoning task. It is a choice-reaction time task, where animals are sequentially presented with a series of shape stimuli, which provide probabilistic evidence. The stimuli consisted of 10 clearly distinguishable shapes (∼ 1.2 degrees of visual angle, dva).

Prior to training, half of the shapes were assigned as evidence in favor of left choices, and the remaining shapes were assigned as evidence in favor of right choices. Of these 10 shapes, two (circle and square) were chosen to act as ‘trump’ shapes: which conferred a ∼ 100 percent likelihood that a saccade to their corresponding targets would result in a reward. The remaining eight shapes were assigned a probabilistic weight, expressed as a log-likelihood ratio, that determined how strongly the shape provided evidence for reward after selecting the corresponding target.

For monkey T positively weighted shapes favored the target presented in the left hemifield and for monkey S these shapes favored the target presented in the right hemifield. For monkey S and monkey T, the weights were [10, 0.9, 0.7, 0.5, 0.3, −0.3, −0.5, −0.7, −0.9, −10] and [10, 0.7, 0.5,0.3, 0.15, −0.15, −0.3, −0.5, −0.7, −10] respectively. The weights for monkey T were reduced compared to monkey S in an attempt to counteract monkey T’s tendency to respond impulsively (initiating saccades after only 1 or 2 shapes), although it had little, if any effect on the animal’s impulsivity.

At the beginning of each trial, 20 shapes were sampled randomly from the distribution associated with the correct target chosen for that trial (Fig. 1). These shapes were stored in an array to be presented consecutively as the trial progressed. As a result, on a small number of trials, the rewarded target could be the right target, while the overall evidence across the 20 shapes favored the left target. In such cases, choosing the left target would be unrewarded.

To start a trial, monkeys were required to direct their gaze to a central fixation point (white annulus, 0.1 dva inner diameter, 0.14 dva outer diameter) and retain fixation for around 600 ms, at which point the two targets (orange and blue annuli, 0.2 dva inner diameter, 0.3 dva outer diameter) would appear at either side of the fixation spot. The position of targets was fixed, with the orange target always appearing to the left and the blue target always to the right side of the fixation point. The first shape cues appeared 400 ms following target presentations. Shape cues were presented sequentially, each for 260 ms, and appeared randomly at one of four locations along an invisible 2 dva x 2 dva grid centered on the fixation spot (Fig. 1), with the exception that a shape never occurred in the same location as the preceding shape. There were 15 ms gaps between shapes. Animals were free to make saccades at any time they felt they had viewed a sufficient number of shapes, that is, accumulated sufficient evidence as to which target was most likely to be rewarded. Trials would be timed out if no decisions were made after the presentation of 20 shapes maximum.

Reaction time (RT) in this task was calculated as the time taken to initiate a saccade from the time of the first shape onset. For saccadic eye movements to be considered as a ‘choice’ to either of the targets, the monkeys’ eye position had to deviate from the fixation window for more than 100 ms, at which point the current shape disappeared and no further shapes were shown. Monkeys then had a maximum of 300 ms to saccade to their selected target and were required to maintain their gaze for a minimum of 350 ms in an invisible 5 dva x 5 dva grid that was centered on the respective targets. A saccade to the rewarded target resulted in a liquid juice reward. If RT was greater than 2,500 ms then the reward delivery was immediate. If RT was less than 2,500 ms the delivery time of the reward was delayed, such that the earliest reward delivery could be at 2,500 ms after first shape onset. This condition was implemented to counteract the monkeys’ natural tendency to respond quickly and was used in the previously published version of this task, along with other reaction time studies using macaques^39^. Trials were separated by a 500 ms inter-trial interval for both monkeys.

### Behavioral models

We fitted the monkeys’ behavior with models that predict the probability of choosing the right target based on the cumulative evidence gathered by observing a sequence of *N* shapes on each trial. In these models, the probability of choosing the right target was a logistic function of the cumulative evidence (Eq. 2). We considered two models—the perfect evidence accumulation model and the working memory limitations (WML) model—which differed in how they transformed the sequence of observed shapes into the cumulative evidence.

The perfect evidence accumulation model assumes that evidence from all shapes sampled before choice remains available to guide the decision. In this model, the cumulative evidence used to compute the probability of choosing the right target is the sum of the subjective weights of all shapes sampled before choice:

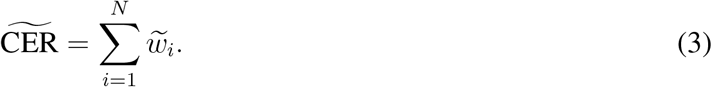

Here *N* is the number of observed shapes, and *w_i_*are their subjective weights, which are the fitted model

The WML model incorporates working memory limitations into the evidence accumulation process, including a memory decay, primacy, recency, and priming. In the WML model, the cumulative evidence is computed as

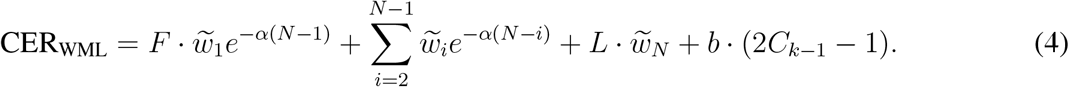

Here, the exponential scaling factors implement memory decay at a constant rate *α*, so that shapes observed earlier in the sequence contribute less to cumulative evidence. The scaling factor *F* implements primacy by multiplying the subjective weight of the first observed shape before it is summed into cumulative evidence. The scaling factor *L* implements recency by multiplying the subjective weight of the last observed shape. The priming term implements a bias toward repeating the previous choice, where *b* controls the strength of the priming effect and *C_k_*_−1_ represents the choice on the previous trial. *C_k_*_−1_ = 1 if the monkey chose the right target on the previous trial and *C_k_*_−1_ = 0 if it chose the left target. The model includes fourteen fitted parameters: 10 subjective weights {*w̃_i_*}, *α*, *F*, *L*, and *b*.

### Fitting behavioral model parameters

We fitted the models by minimizing a loss function defined as the negative log-likelihood of monkey choices for the corresponding shape sequences in the dataset:

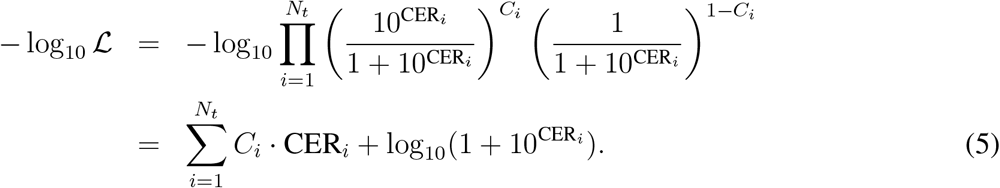

Here, *C_i_*= 1 if the monkey chose the right target on trial *i*, and *C_i_* = 0 if it chose the left target. *N_t_* is the number of fitted trials. For each trial *i*, cumulative evidence CER*_i_* is computed from the sequence of observed shapes as [inline] (Eq. 3) for the perfect evidence accumulation model and as CER_WML_ (Eq. 4) for the WML model. To determine the unique contribution of each factor—memory decay, primacy, recency, and priming—we systematically removed each factor from the WML model and fitted the resulting reduced model to the monkeys’ behavior.

We minimized the loss Eq. 5 using unconstrained simplex search method (fminsearch in MATLAB). When fitting models with memory decay, we used constrained minimization with the interior-point method (fmincon in MATLAB), restricting the decay parameter *α* to the range [0, 1].

We used bootstrapping to estimate errors. For the analysis in Fig. 5, we performed 100 bootstrap iterations. In each iteration, we resampled trials with replacement, using a sample size equal to the total number of trials.

### Mixed-effects model of the decision threshold

Several factors can influence the variability in the decision threshold, defined as the amount of subjective cumulative evidence required to commit to a choice. First, the decision threshold may change with the number of observed shapes. For example, the decision threshold may decrease as the number of sampled shapes grows, reflecting urgency to commit to a choice. We modeled this effect with a polynomial function of the number of sampled shapes, *n*. Second, the decision threshold may change over the course of a session, reflecting gradual shifts in the animal’s motivation, engagement, or impulsivity. We modeled this effect with a polynomial function of the time elapsed since session onset, *t*. Finally, the decision threshold may vary across sessions. We modeled this effect with a random intercept (1|session), representing a session-specific baseline for the decision threshold.

To disentangle these different sources of variability, we fitted the subjective cumulative evidence at decision time with a mixed-effects model:

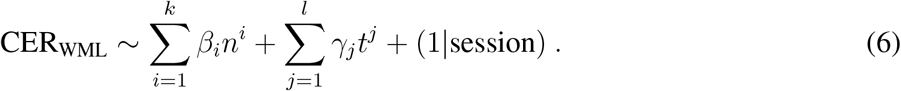

Here *k* and *l* are the degrees of fitted polynomials, and coefficients *β_i_* and *γ_j_* are free parameters that control the shape of these polynomial functions. We fitted the model parameters using the fitlme function in MATLAB. We performed a grid search over integer values of *k* and *l* in the range [1, 5] and selected the highest polynomial degrees for which the corresponding coefficients, *β_k_* and *γ_l_*, remained significant. This procedure selected polynomial degrees of *k* = 4 and *l* = 2 for monkey S, and *k* = 4 and *l* = 1 for monkey T.

### Neural circuit model

The network consists of two integrator units, representing two neural populations that accumulate evidence for the left (L) and right (R) choices. The integrator units receive inputs from 10 input units, each activated by a specific shape. The firing rate of input unit *i* is set to *r_i_* = 1 while shape *i* is present on the screen and to zero otherwise. The firing-rate dynamics of the integrator units are governed by:

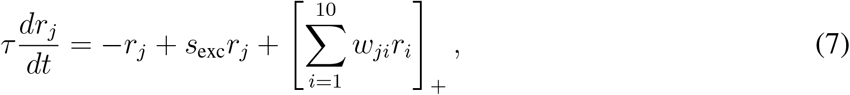

where ∀*j* ∈ {L, R}. The time constant *τ* is 10 ms. We optimized the self-excitation parameter *s*_exc_ and connection weights *w_ji_* to fit the model to the behavior for each monkey separately.

We fitted the model by minimizing the same negative log-likelihood of monkey choices as for the behavioral models (Eq. 5):

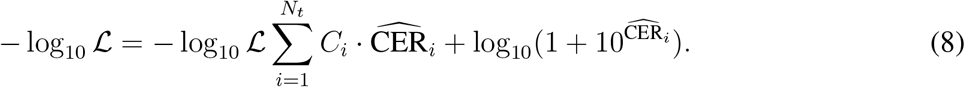

For the neural circuit model, the cumulative evidence is defined as 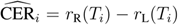, the firing-rate difference between the right and left integrator units at reaction time *T_i_* on trial *i*. The choice variable *C_i_* = 1 if the monkey chose the right target on trial *i*, and *C_i_* = 0 if it chose the left target. *N_t_*is the number of fitted trials. We used all trials from all sessions for model training. Since the dataset was large relative to the number of fitted parameters (∼ 100, 000 trials and 12 parameters), we did not use a separate validation set. Model parameters were optimized by gradient descent using the Adam optimizer with a learning rate of 0.02, minibatches of 128 trials, and a fixed training duration of 10 epochs. L2 regularization was applied through Adam weight decay of 10^−3^. The number of epochs was fixed a priori and was not selected using early stopping or validation-set performance. To assess robustness to initialization, we repeated the fitting procedure across 30 different initializations.

### Drug preparation and administration

We used SKF81297 (a selective D1 receptor agonist) at four different doses. There were 0.02 (Monkey S), 0.05 (Monkey T), 0.1, 0.2, 0.3 mg/kg. SCH23390 (a selective D1 receptor antagonist) was also used at four different doses at 0.05, 0.01, 0.02, 0.03 mg/kg. Sterile saline (Sodium Chloride 0.9 percent) was used as a control. Both dopaminergic (DA-ergic) agents were obtained from Bio-Techne ltd. Drug solutions for injections were made either on the day of the experiment or at the start of an experimental week. All drugs were prepared under aseptic conditions and refrigerated within sterile universal tubes, until a minimum of an hour prior to administration. This ensures that drug solutions are at room temperature when injected. Drugs were kept refrigerated for up to 4 days. For initial experimental sessions, drugs were administered systemically intramuscularly (IM) into the monkeys’ thighs. For the later sessions, we used subcutaneous (SC) injection, as this reduced disturbance and stress for the animal. Here a butterfly cannula was placed and secured into a skin fold at the monkey’s neck prior to the experiment and the injection could be done with minimal disturbance after the pre-injection data had been acquired, using an automated injection pump triggered remotely (injection rate 0.2 ml/min).

Experimental sessions were completed over a series of weeks, whereby either, the agonist (DA+) or antagonist (DA-) drug would be chosen by the experimenter as the chosen agent of that week. This drug would then be administered on alternate days, with saline being the administered agent on the intermittent days. This procedure ensured we provided a significant ‘wash-out’ period for the DA-ergic agent, but also avoided any cumulative effects over a series of days. The weeks were structured such that, for one week, Saline would be the administered agent on experimental Day 1, followed by the DA-ergic agent on Day 2. The following week this order was reversed so that the DA-ergic agent was administered on Day 1 and the Saline on Day 2. Within a single week, the investigated drug doses were chosen by the experimenter. In the event that two different doses were chosen within the same week, the lower dose was administered on either Day 1 or Day 2 and the higher dose on Day 3 or Day 4.

To investigate changes in the monkeys’ behavior following the administration of DA-ergic drugs on any given experimental day, we obtained both a series of PRE and POST-Injection trials. This allowed us to identify any changes in POST-injection behavior compared to the respective baseline behavior observed in the PRE-injection trials on a daily basis. Order reversal was not possible as sessions were not long enough to ensure that a complete washout was achieved within a day. The minimum number of shape trials required for both PRE and POST-injection trials was set to 180 for both subjects. For PRE-injection trials, after collection of ∼180 trials, the task was manually terminated by the experimenter, after which the drug/control was administered to the monkey. The monkeys then began the POST-injection trials immediately and were allowed to work for as long as they appeared motivated to do so, which typically exceeded the 180-trial benchmark.

### Fitting behavioral model parameters across drug conditions

We fitted the WML model with all 14 parameters treated as free parameters. To improve parameter-estimation precision, we pooled trials across different doses of the same drug. For each drug condition, we performed 100 bootstrap iterations, randomly sampling 4,000 trials without replacement in each iteration.

### Mixed-effects model of effect of drug on number of shapes

The number of shapes that a monkey observes during the task may be influenced by several factors, including the session, the number of trials completed (e.g., towards the end of the session, the monkey may observe fewer shapes due to fatigue or decreased motivation), the type of injection administered, and the effect of the drug. To isolate the drug effect, we used a mixed-effects model where the number of shapes *N* was modeled as:

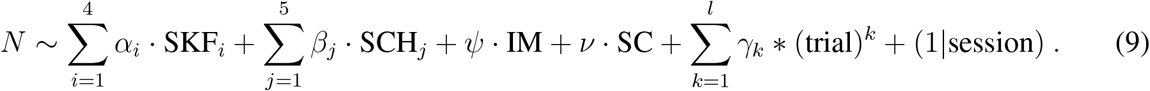

Here, (1|session) is a random effect, allowing each session to have its own baseline shift in the number of shapes, which accounts for session-specific variability. The terms *α_i_* and *β_j_* capture the effects of the different doses of the SKF and SCH drugs, respectively. Specifically, SKF*_i_* = 1 only if dosage *i* of SKF is applied in that trial, and similarly for SCH*_j_*. The variables IM and SC are indicator variables for the type of injection: intramuscular (IM) or subcutaneous (SC). These terms equal 1 only for the trials after the injection and 0 otherwise. We include these factors because the type of injection may affect the monkey’s behavior, potentially influencing the number of shapes observed. To model the relationship between the number of shapes and the number of trials elapsed, we use a polynomial function. We incrementally increase the order *k* of the polynomial until the coefficients *γ_k_* are no longer statistically significant. Using the fitlme function in MATLAB, we determined that polynomial terms beyond *k* = 3 were non-significant in our model.

## Acknowledgements

This work was supported by the Swartz Foundation (C.A. and C.L.), the National Institutes of Health (NIH) grant R01 EB026949 (T.A.E., C.A., C.L.), NIH grant RF1DA055666 (T.A.E., C.A., C.L.), NIH grant S10OD028632-01 (T.A.E., C.A.), Alfred P. Sloan Foundation Research Fellowship (T.A.E.), Wellcome Trust [093104] (A.T.), MRC [MR/P013031/1] (A.T.), and BBSRC [BB/W006758/1] (A.T).

## Author contributions

J.v.K., M.S., A.G., and A.T. conducted the experiments. C.A., T.A.E., and A.T. developed the behavioral models. C.A. developed the code for fitting the behavioral models and analyzed the experimental data. C.A., C.L., and T.A.E. developed the neural circuit model, and C.A. and C.L. developed the associated code. C.A., A.T., and T.A.E. wrote the manuscript.

## Data availability

Data will be made available upon publication of the manuscript at g-node.org.

## Code availability

The source code to reproduce the results of this study will be made available on GitHub upon publication.

