## Supplementary Information for "Working memory limitations and dopamine modulation in probabilistic reasoning"

August 13, 2026

### 1 Supplementary Notes

#### 1.1 Estimating the effective evidence accumulation window

To quantify each monkey's evidence-sampling strategy, we first determined which shapes were presented early enough before saccade onset to influence the choice. This distinction was necessary because shapes continued to appear until the monkey made a saccade, whereas motor planning and execution require time. Consequently, shapes displayed immediately before the saccade might not have been incorporated into the decision. To identify when decision formation effectively terminated, we fitted a model to the monkeys' choices. In this model, the probability of choosing the right target depended on the cumulative evidence favoring that target, defined as CER $= \sum_i g(t_i)w_i$. Here, $g(t_i)$ is the gain, or relative influence, of the shape presented at time $t_i$ relative to saccade onset, with $t_i = 0$ corresponding to the time of the saccade. We assumed that all shapes

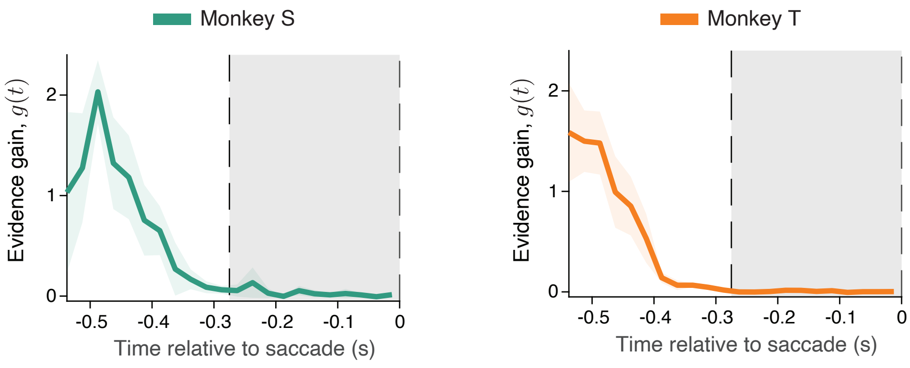

**Supplementary Figure 1. Influence of shapes presented before saccade onset on the monkeys' choices.** The gain $g(t_i)$ governing the contribution to CE of the shape presented at time $t_i$ before a saccade onset (left panel: monkey S, right panel: monkey T). Gray shading indicates the 260 ms interval before saccade onset, corresponding to the maximum duration for which the final shape remained on the screen.

presented more than 550 ms before the saccade—equivalent to two shape-presentation intervals—had unit gain, such that $g(t_i) = 1$ for $t_i < -550$ ms. Shapes presented closer to the saccade were allowed to have different gains. We divided the interval $-550 \leqslant t_i < 0$ ms into 25-ms bins and fitted $g(t_i)$ in each bin as an independent free parameter. For both monkeys, shapes present on the screen during approximately the final 400 ms before the saccade had little to no effect on the monkeys' choices (Supplementary Fig. 1). Therefore, in all further analyses, we do not include the final shape presented before each saccade, which provides a more accurate estimate for the number of shapes each animal used for making a choice.

#### 1.2 Ideal Bayesian observer

To establish a normative benchmark for evaluating the monkeys' choices, we derived the decision strategy of an ideal Bayesian observer in the probabilistic reasoning task. On every trial, each shape is sampled with a different probability depending on whether the right or left target is rewarded (Fig. 1b). Thus, each observed shape provides probabilistic evidence about which target is rewarded. In a Bayesian framework, this evidence is given by the log-likelihood ratio of observing that shape when the right target is rewarded versus when the left target is rewarded. This log-likelihood ratio defines the weight $w_i$ of shape $i$:

$$w_i = \log_{10} \frac{\mathbb{P}(\mathbf{Target} = \text{right}|\mathbf{Shape} = i)}{\mathbb{P}(\mathbf{Target} = \text{left}|\mathbf{Shape} = i)} = \log_{10} \frac{\mathbb{P}(\mathbf{Shape} = i|\mathbf{Target} = \text{right})}{\mathbb{P}(\mathbf{Shape} = i|\mathbf{Target} = \text{left})},$$

where the second equality assumes equal prior probabilities for the two targets.

The rewarded target is chosen at the start of each trial, and then each shape is drawn independently from the respective distribution. Since successive shapes are drawn independently conditional on the rewarded target, their log-likelihood ratios add across the sequence. Thus, after observing $n$ shapes, the cumulative evidence for the right target is:

$$\begin{aligned}\text{CE} &= \log_{10}\frac{\mathbb{P}(\textbf{Target}=\text{right}|\textbf{Shape}=s_1s_2\cdots s_n)}{\mathbb{P}(\textbf{Target}=\text{left}|\textbf{Shape}=s_1s_2\cdots s_n)}\\ &= \log_{10}\frac{\mathbb{P}(\textbf{Shape}=s_1s_2\cdots s_n|\textbf{Target}=\text{right})}{\mathbb{P}(\textbf{Shape}=s_1s_2\cdots s_n|\textbf{Target}=\text{left})}\\ &= \sum_{i=1}^{n}\log_{10}\frac{\mathbb{P}(\textbf{Shape}=s_i|\textbf{Target}=\text{right})}{\mathbb{P}(\textbf{Shape}=s_i|\textbf{Target}=\text{left})}=\sum_{i=1}^{n}w_{s_i}.\end{aligned}$$

Since $\mathbb{P}(\textbf{Target}=\text{left}|\textbf{Shape}=s_1s_2\cdots s_n)=1-\mathbb{P}(\textbf{Target}=\text{right}|\textbf{Shape}=s_1s_2\cdots s_n)$, the probability of the right target being rewarded after observing the first $n$ shapes equals

$$\mathbb{P}(\textbf{Target}=\text{right}|\textbf{Shape}=s_1s_2\cdots s_n)=\frac{10^{\sum_{i=1}^{n}w_{s_i}}}{1+10^{\sum_{i=1}^{n}w_{s_i}}}.$$

This posterior probability represents the belief that an ideal Bayesian observer assigns to the right target being rewarded after observing the sequence. This Bayesian posterior probability serves as the probability-matching benchmark for monkey behavior (Fig. 2d).

#### 1.3 Trade-off in continued evidence sampling under memory decay

working memory decay alters the criterion for terminating evidence sampling because the benefit of acquiring additional evidence decreases as the sequence progresses. Although each newly presented shape is drawn from the same target-conditioned distribution and therefore provides the same expected amount of evidence, sampling it comes at the cost of decay in previously accumulated evidence. We used the WML model to quantify this trade-off by calculating the expected marginal gain in subjective cumulative evidence from observing one additional shape and comparing this benefit with the time cost of continued sampling.

In the WML model, the subjective cumulative evidence after observing $n$ shapes is

$$\text{CER}_{\text{WML}}(n)=F\mathbf{X}_1e^{-\alpha(n-1)}+\sum_{k=2}^{n-1}\mathbf{X}_ke^{-\alpha(n-k)}+L\mathbf{X}_n+b\mathbf{Y}. \tag{1}$$

Here $X_k$ is the subjective evidence weight associated with the $k$th observed shape, and $Y$ is a random variable that models the priming effect, with $Y=1$ if the monkey's previous choice was right and $Y=-1$ if it was left. The parameter $\alpha$ represents the memory decay rate, and the parameters $F$, $L$, and $b$ are associated with the primacy, recency, and priming effects, respectively. Since the shapes are generated independently from the same target-conditioned distribution within a trial, the expected value of $\mathbf{X}_k$, given that the target is "right", is constant for all $k$. We denote this expected value by $Q$, that is, $\mathbf{E}[X_k|\textbf{Target}=\text{right}]=Q$. Then, the mean of the subjective cumulative evidence after observing $n$ shapes, given that the correct target is "right", is

$$\mathbf{E}[\text{CER}_{\text{WML}}(n)|\textbf{Target}=\text{right}]=FQe^{-\alpha(n-1)}+\sum_{k=2}^{n-1}Qe^{-\alpha(n-k)}+LQ+b\mathbf{E}[Y]. \tag{2}$$

We next examine how the expected cumulative evidence in Eq. 2 grows with the number of sampled shapes, which reveals the trade-off associated with continued evidence sampling. From

Eq. 2, we see that, excluding the priming bias, the expected subjective evidence accumulated from a shape sequence approaches the asymptotic limit $Q(L + \frac{e^{-\alpha}}{1-e^{-\alpha}})$ as the number of sampled shapes increases ($n \to \infty$). Thus, the expected evidence accumulated from the shape sequence is bounded and cannot increase indefinitely. We define the expected marginal gain in subjective cumulative evidence from sampling the $n$th shape as

$$\Delta_{\text{evid}}(n) = \mathbf{E}[\text{CER}_{\text{WML}}(n)|\mathbf{Target} = \text{right}] - \mathbf{E}[\text{CER}_{\text{WML}}(n-1)|\mathbf{Target} = \text{right}].$$

With this definition, we obtain

$$\Delta_{\text{evid}}(n) = Q(1 - F(1 - e^{-\alpha})) \cdot e^{-\alpha(n-2)}.$$

Thus, the expected marginal gain from sampling an additional shape decreases exponentially with sequence length $n$, since newly acquired evidence is increasingly offset by the decay of previously accumulated evidence. Assuming that sampling each additional shape incurs a constant time cost, $c_{\text{time}}$, independent of sequence length, the expected net gain from extending the sequence from $n-1$ to $n$ shapes is $\Delta_{\text{total}}(n) = \Delta_{\text{evid}}(n) - c_{\text{time}}$. Since $\Delta_{\text{evid}}(n)$ decreases with sequence length while $c_{\text{time}}$ remains constant, continued sampling eventually ceases to be beneficial. Specifically, once $\Delta_{\text{evid}}(n) < c_{\text{time}}$, the expected cost of sampling an additional shape exceeds its expected gain in subjective evidence, favoring termination of evidence sampling. Thus, working memory decay reduces the benefit of continued sampling as sequences lengthen. Moreover, all else being equal, faster memory decay causes the marginal gain in evidence to fall below the time cost sooner, predicting termination of evidence sampling after fewer shapes.

#### 1.4 Dopamine effects on primacy and recency

We examined the effects of D1R manipulation on the serial-position biases captured by the primacy and recency parameters in the WML model. D1R antagonist (SCH) did not significantly alter primacy in either monkey, whereas D1R agonist (SKF) had opposite effects on primacy in the two animals (Supplementary Fig. 2). For recency, D1R agonist reduced the contribution of the last shape in both monkeys, whereas D1R antagonist affected the last-shape contribution differently across the two animals. These heterogeneous effects may reflect differences in the animals' baseline positions along the inverted-U-shaped relationship between D1R signaling and working memory function.

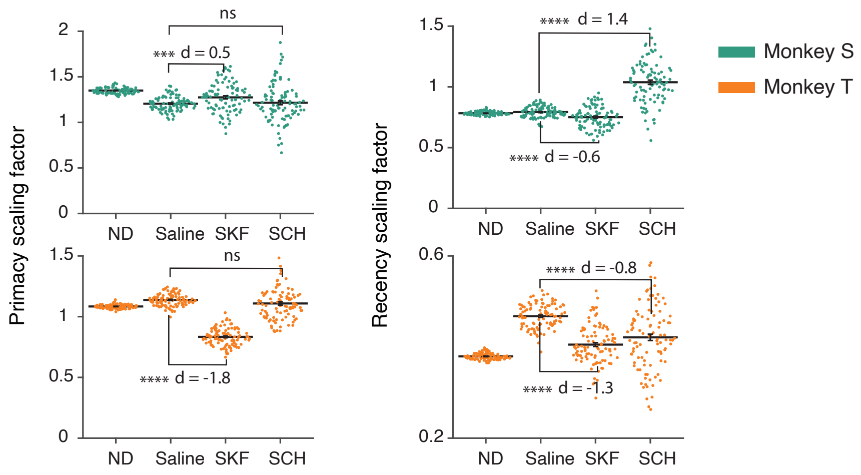

**Supplementary Figure 2. Dopamine modulation of primacy and recency.** The effect of the dopamine agonist (SKF) and the antagonist (SCH) drugs on the contribution of first (primacy) and last shape (recency) to the cumulative evidence. The p-values are from two-tailed t-test ($^{***}p < 10^{-3}$, $^{****}p < 10^{-10}$).
